# Genome-wide kinetic profiling of pre-mRNA 3’ end cleavage

**DOI:** 10.1101/2023.06.21.545926

**Authors:** Leslie Torres Ulloa, Ezequiel Calvo-Roitberg, Athma A. Pai

## Abstract

Cleavage and polyadenylation is necessary for the formation of mature mRNA molecules. The rate at which this process occurs can determine the temporal availability of mRNA for subsequent function throughout the cell and is likely tightly regulated. Despite advances in high-throughput approaches for global kinetic profiling of RNA maturation, genome-wide 3’ end cleavage rates have never been measured. Here, we describe a novel approach to estimate the rates of cleavage, using metabolic labeling of nascent RNA, high-throughput sequencing, and mathematical modeling. Using in-silico simulations of nascent RNA-seq data, we show that our approach can accurately and precisely estimate cleavage half-lives for both constitutive and alternative sites. We find that 3’ end cleavage is fast on average, with half-lives under a minute, but highly variable across individual sites. Rapid cleavage is promoted by the presence of canonical sequence elements and an increased density of polyadenylation signals near a cleavage site. Finally, we find that cleavage rates are associated with the localization of RNA Polymerase II at the end of a gene and faster cleavage leads to quicker degradation of downstream read-through RNA. Our findings shed light on the features important for efficient 3’ end cleavage and the regulation of transcription termination.

## INTRODUCTION

Cleavage at the 3’ end of a pre-mRNA molecule is a quintessential part of mRNA maturation. This step determines the composition of the 3’ untranslated region (UTR) and allows the release of mRNA from the active polymerase and chromatin environment (1). Cleavage is initiated by the recognition of a polyadenylation site (PAS) (2), which stimulates the cleavage and polyadenylation machinery to perform endonucleolytic cleavage downstream of the PAS while RNA Polymerase II (RNAPII) continues transcribing until other processes terminate transcription and degrade the read-through transcript (3). An mRNA molecule is fully mature only after the addition of a tail of non-templated adenines to the 3’ end (polyA tail). The coupling of cleavage and polyadenylation protects mRNAs - together with the 5’ N7-methylguanosine cap - from premature degradation during nuclear export to the cytoplasm (4–6). The spatio-temporal coordination of these processes is likely heavily regulated to ensure productive mRNA expression.

Eukaryotic mRNAs often contain many PASs, with extensive alternative polyA site usage leading to isoforms with alternative 3’ UTRs. 3’ UTR composition can influence polyA tail length (7, 8), export efficiency (8), mRNA stability(9), mRNA localization (10, 11), and translation efficiency(12, 13). The usage of alternative polyA sites within upstream coding exons or introns can create truncated transcripts that result in altered protein composition or are targeted for degradation. Alternative PAS usage has been implicated in triggering many disease mechanisms (2, 14), developmental or differentiation states (15, 16), or cellular responses to stressors such as immune stimuli or heat shock (17–19). The recognition of PASs is regulated through a combination of local *cis-*regulatory sequences and global changes in the availability of necessary *trans-*regulatory factors, and transcription elongation rates (2, 20). The relative rates at which polyA sites are recognized within or across genes likely have a direct impact on the relative expression of different isoforms.

The development of high-throughput sequencing approaches to systematically capture and analyze nascent pre-mRNA molecules has opened the door for genome-wide kinetic profiling of RNA maturation processes (21, 22). Many steps of RNA maturation - including mRNA splicing and cleavage - are thought to occur primarily co-transcriptionally (20), with substantial interplay between the efficiency and choices involved in each of these processes. Preliminary evidence suggests that intronic splice site and PAS choices are determined by kinetic parameters, such that sequence features or gene architecture might promote faster intron excision or transcript cleavage(23, 24). While these hypotheses are being increasingly tested and molecularly characterized for mRNA splicing (20), we still have very little understanding of the kinetic regulation of PAS choice and mRNA cleavage that determines the composition of the 3’ ends of mRNAs.

Previous studies have suggested that CPA is likely to occur relatively rapidly after RNA synthesis and can be temporally regulated (25–27). An early molecular study measured the addition of radiolabeled adenosines to the 3’ ends of pre-mRNAs synthesized with radiolabeled uridines and observed that polyadenylation (and thus cleavage) likely occurred within 1 minute of mRNA biogenesis in CHO and HeLa cells (25). However, this study only measured global RNA polyadenylation, with no ability to measure variation in the rates of cleavage across genes or alternative sites. More recently, insights from high-throughput sequencing of chromatin associated RNA suggested that variation in splicing and 3’ end kinetics (rather than transcription initiation rates) across genes controlled the timing of gene expression for multiple sets of immune genes. Similarly, imaging studies have suggested that there is kinetic competition between splicing and cleavage in the process of mRNA maturation and that the temporal balance between these mechanisms can be regulated (27). Together, these studies highlighted the need to more precisely estimate site-specific rates of cleavage and polyadenylation that govern the timing that mature mRNAs are available for downstream processes.

Here, we use nascent RNA high-throughput sequencing and mathematical modeling to develop a method for estimating site-specific 3’ cleavage rates with temporal precision. Our approach is based on our previous method for estimating intron-specific rates of mRNA splicing, which models the probability of observing splicing-informative junction reads across time since synthesis of the pre-mRNA molecule. To use this framework to estimate rates of cleavage, we first use simulations to (1) characterize reads informative about uncleaved and cleaved molecules, (2) derive a model that measures cleavage half-lives given the probability of observing a ratio of uncleaved to cleaved molecules given time since pre-mRNA synthesis, and (3) extend this model to estimate the rates of cleavage at alternative cleavage sites within a gene. Applying this model to nascent RNA data from *Drosophila melanogaster* S2 cells, we estimate 3’ cleavage half-lives for approximately 3, 000 cleavage sites, identify sequence features associated with the rates of cleavage, and characterize RNA Polymerase II (RNAPII) dynamics around sites with variable cleavage half-lives.

## METHODS

### Processing 4sU-seq data

4sU-seq data from *Drosophila melanogaster* S2 cells were obtained from Pai *et al.* 2017 (24). This dataset consists of three independent replicates each of 4sU-seq data after 5, 10, and 20 minutes of labeling with 500 micrograms of 4sU as described in Pai *et al.* (24) and two replicates of total RNA. These data include 51nt paired-end reads and 100nt reads for the total RNA-seq samples, all sequenced on an Illumina HiSeq platform. Fastq files were downloaded from GEO (GSE93763) and mapped to the dm6 reference genome (Flybase release 6.28) scaffolded with BGDP6.28.99 annotations using STAR (version 2.7.0e) (56) with options: --outSAMtype BAM SortedByCoordinate --outSAMstrandField None --outSAMattributes All --alignIntronMin 25 --alignIntronMax 300000]. Gene expression levels were calculated for the total RNA-seq data with Salmon (57) using the dm6 6.28 reference transcriptome. Gene level TPMs were obtained by summing all the isoform-level TPMs for a given gene.

### Identification and estimation of cleavage site usage

To identify cleavage sites that are used in *Drosophila melanogaster* S2 cells, we used a published 3p-seq data from S2 cells (58) and retained cleavage sites in the dm6 6.28 annotations that overlapped (within +/- 25nt) with 3p-seq peaks after lifting over the curated table from dm3 coordinates.

Alternative usage of cleavage sites was measured using a combination of the LABRAT and QAPA pipelines to measure proportional usage of cleavage sites. We used LABRAT (version 0.2.2) (59) to filter annotated transcription end sites and calculate the expression of terminal fragments in total RNA-seq data. We then used the Poly(A) Usage (PAU) metric introduced by QAPA (60) to estimate a site-specific PAU value, which represents the proportion of cleavage site usage relative to the usage of all other cleavage sites in a gene.

### Estimating 3’ end cleavage rates with mathematical modeling of informative read ratios

We based our model for estimating the rate of 3’ end cleavage on our previous model for estimating the rate of mRNA splicing (24). The progressive labeling design implemented in the 4sU-seq experiments enables the capture of RNAs at different stages of processing, and yields three populations of transcripts: (1) transcripts not yet transcribed past the cleavage site; (2) transcripts that are fully transcribed but still uncleaved; (3) transcripts that are fully transcribed and also cleaved. As described in Pai *et al.*, this progressive metabolic labeling data, we model the proportion of reads belonging to either cleaved or uncleaved transcripts as a function of the distance traveled by RNA Polymerase II and the probability of observing either cleavage state at a given polymerase endpoint (24). This is distinct from parameterizing the model by timepoints, since conceptualizing probability in terms of polymerase distances allows the modeling of a range of transcript lifetimes within the same labeling timepoint.

To identify reads that are directly informative about 3’ end cleavage state, we define two kinds of informative regions: (1) overlapping a cleavage site, where reads can only come from uncleaved transcripts, and (2) two insert lengths upstream of the cleavage site, where reads can come from both uncleaved and cleaved transcripts and are informative about the total number of transcripts that can be cleaved at this site. We use this second region to estimate the expected number of reads coming from cleaved transcripts by subtracting the reads overlapping the cleavage site and the number of reads estimated to be derived from transcripts have transcribed past the total region but have not yet reached the cleavage site (**FIGURE 1C; FIGURE S1**). We use these empirical estimates of uncleaved and cleaved transcripts to mathematically model the expected 3’ end cleavage half-life across a range of transcript lifetimes. We use the same mathematical expression described to estimate splicing half lifes in Pai *et al.* (24), with three important changes: (1) the empirical ratio is of uncleaved/cleaved transcripts, rather than unspliced/spliced transcripts, (2) only transcripts that have transcribed past the cleavage site are considered, and (3) the range of polymerase endpoints is redefined to consider the region that is informative about cleavage, rather than splicing.

**Figure 1.**
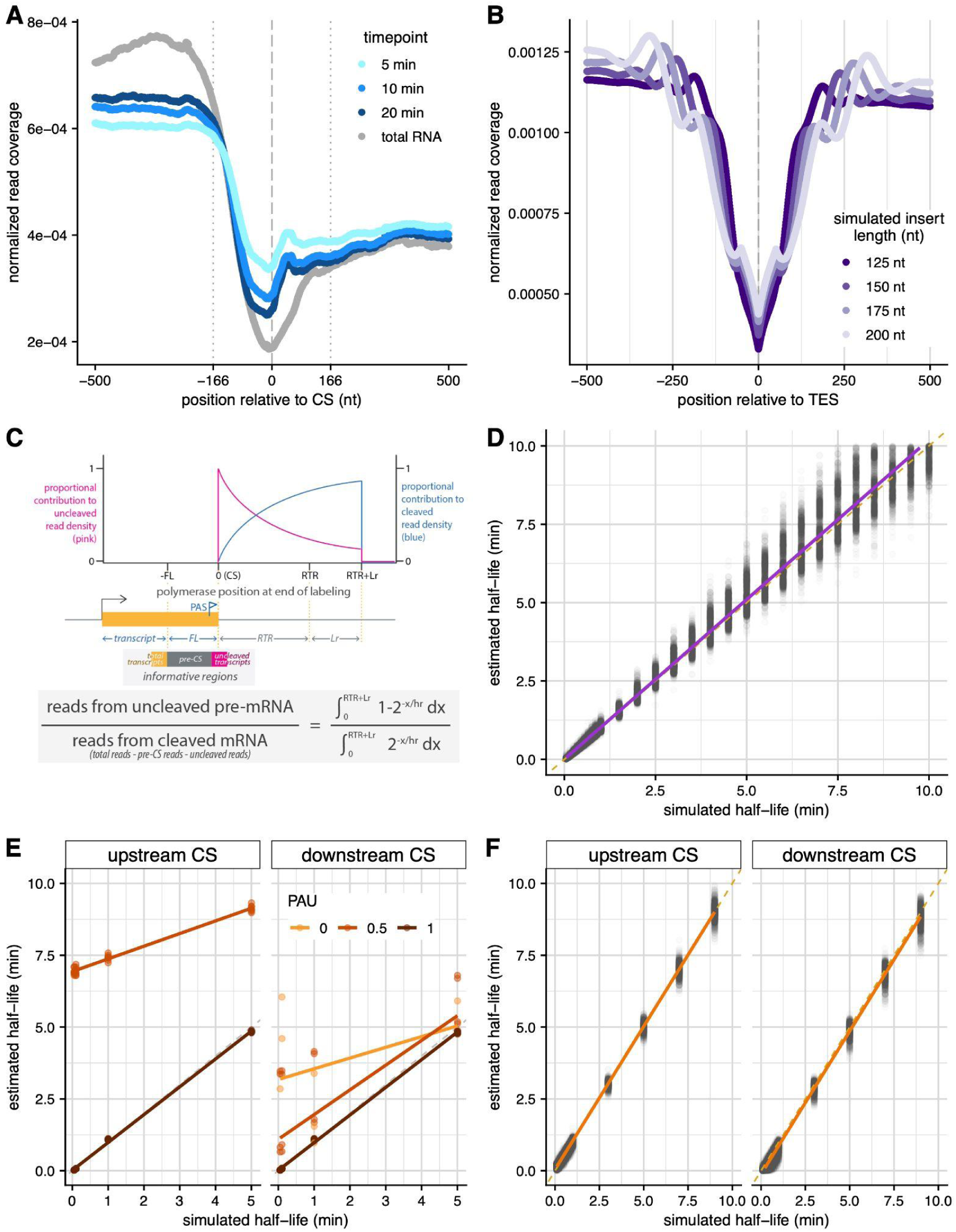
Developing a mathematical model to estimate site-specific rates of 3’ end cleavage. **(A)** Meta-gene plot showing average normalized 4sU-seq read coverage (*y-axis*) in a 1000nt region centered on annotated 3’ end cleavage sites (*x-axis*), separated by 4sU labeling timepoints (*blue lines*) and steady-state total RNA data (*grey line*). The dashed line indicates the cleavage site (coordinate 0), while dotted lines indicate the library fragment length. **(B)** Meta-gene plot showing the average simulated read coverage (*y-axis*) in a 1000nt region centered on a simulated 3’ end cleavage site (*x-axis*), separated by variable simulated library insert lengths (*purple lines*). **(C)** Schematic of the expected read distribution resulting from progressive labeling with 4sU, as described in Pai *et al.*. The probability of sampling uncleaved (*pink*) and cleaved (*blue*) transcripts is dependent on the cleavage half-life and polymerase position at the completion of the labeling period. Reads overlapping the cleavage site can be used to approximate uncleaved transcripts, while reads located in a region upstream of the cleavage site (defined by the library fragment length) are informative of the total number of transcripts and can be used to calculate the approximate number of cleaved transcripts. **(D)** 3’ end cleavage half-lives estimated by the mathematical model (*y-axis*) versus half-lives used to simulate read data (*x-axis*). Dotted yellow line indicates y=x line of perfect correlation, while purple line indicates linear fit to the data. **(E)** Cleavage half-lives estimated by the approach for constitutive cleavage sites (*y-axis*) for simulated alternative cleavage sites (*panels*) versus half-lives used to simulate read data (*x-axis*). Dotted grey line indicates y=x line of perfect correlation, while colored lines indicate linear fits for sites with variable alternative usage. **(F)** Cleavage half-lives for alternative cleavage sites estimated after proportionally correcting read counts by relative usage (*y-axis*) versus half-lives used to simulate read data (*x-axis*). Shown are estimates for all upstream cleavage sites (*left*) and downstream cleavage sites with a simulated PAU > 40% (*right*). Dotted orange line indicates y=x line of perfect correlation, while orange line indicates linear fit to the data.

Specifically, considering the cleavage site (*CS*) to be coordinate 0, the range of polymerase endpoints informative for cleavage becomes [*CS*, *RTR*+*Lr*], where *RTR* is the length of the readthrough region, *L* is the labeling time, and *r* is the transcription rate, assumed to be constant (**FIGURE 1C**). Transcripts associated with polymerases located upstream of CS cannot yet be cleaved, and transcripts associated with polymerases located beyond *RTR*+*Lr* cannot have incorporated the metabolic label. Using these new integration limits, the model can be solved for the half-life *h* to obtain an estimate of a 3’ end cleavage half-life. However, since there is no analytical solution for the resulting function, we solve a simplified expression numerically using the *optim* function in R, as described in Pai *et al.* (24). Similar to the model to estimate splicing half-lives (24), this model to estimate 3’ end cleavage half-lives assumes (1) an equal probability of sampling reads from uncleaved and cleaved transcripts, (2) independence between cleavage events, and (3) that all transcripts will eventually be cleaved at the site where cleavage half-life is measured.

#### Estimate 3’ end cleavage half-lives for alternative cleavage sites

While the assumptions described above are valid for constitutively used cleavage sites, they are not all valid for sites that are alternatively used (PAU < 95%). Specifically, there is an intrinsic interdependence between the counts of reads across alternative sites from the same gene since transcripts that are cleaved at a given site must be uncleaved at the other site. This results in the probability of cleavage at a site being dependent on the cleavage status of other sites in the gene. This relationship results in an overcounting of uncleaved reads at sites with PAU < 95% (**FIGURE S3B and C**). Thus, to obtain cleavage rates estimates for alternative sites with 5% < PAU < 95%, we must proportionally allocate informative reads based on the site-specific usage observed in steady-state data (**FIGURE S4**). Broadly, we (1) use site-specific PAU values to weight the observed counts of reads at the total region and each region directly overlapping consecutive cleavage sites, (2) subtract reads that are assumed to originate from transcripts that are not able to be cleaved at a given site, and (3) calculate the expected number of reads from cleaved transcripts by subtracting the adjusted uncleaved read count from the adjusted total read count. To enable these calculations, we count reads from the same region upstream of the first cleavage site (as described above) to obtain the total number of transcripts for a gene. Furthermore, we make a few key assumptions, primarily that there is again a uniform distribution of polymerases throughout the entire 3’ end of the gene and that each pre-mRNA molecule is only cleaved once, after which there is relatively fast degradation of the readthrough fragment.

Specifically, for the initial CS_1_ we perform the following series of calculations:

1. The number of reads from transcripts that can be cleaved at CS_1_ is calculated by weighting the read count from the total region by PAU_1_ and then subtracting the expected proportion of reads arising from polymerases that are within the pre-CS_1_ region.
2. The uncleaved read count from CS_1_ is corrected by subtracting the expected proportion of reads arising from transcripts that will be cleaved at all downstream cleavage sites.

For all downstream expressed cleavage sites *n*, we perform the following series of calculations:

1. The number of reads from transcripts that can be cleaved at CS_n_ is calculated by weighting the read count from the total region by PAU_n_ and then subtracting (a) the expected proportion of reads arising from polymerases that are within the pre-CS_1_ region and (b) the expected proportion of reads arising from polymerases that are within the extended UTR region defined as CS_n_ - CS_1_.
2. The uncleaved read count from CS_n_ is corrected by subtracting the uncleaved read count from all CS_i<n_, after subtracting for the number of those reads that are expected to arise from polymerases that are within the extended UTR region defined as CS_n_ - CS_i<n_ for all CS_i<n_.

### Simulations to assess modeling approaches

We used simulated 4sU-seq data to assess the accuracy of our 3’ end cleavage rate models. To do so, we simulated data for a cleavage site across a range of different biological and technical contexts: gene length *g* (1 - 50kb), readthrough length RTR (1 - 100 kb), expression level in TPM (10 - 250 TPM), labeling periods *L* (5, 10, and 20 min), mean library preparation insert size (125 - 200 nt), mean standard deviation around the insert size (25 nt), and half-lives *h* (0.1 - 30 min). For simulations where we simulated two cleavage sites in the same gene, we also simulated ranges of proportional cleavage site usage (0 - 100% PAU) and distances between cleavage sites *d* (500 - 2000 nt). These parameters recapitulate standard distributions of gene structures, expression values, and experimental conditions. We note that the simulations allow RNAPII to transcribe the full RTR distance, but do not simulate XRN2 cleavage of readthrough transcripts and thus we remove reads from the RTR for kinetic modeling of simulated data.

To generate read data from transcripts, we simulated several steps in nascent RNA transcription, capture, and library preparation as previously described in Pai *et al.* (24), and modified to simulate 3’ end cleavage rather than splicing processes. Most notably, for transcripts with RNAPII endpoints beyond the cleavage site, the cleavage state was probabilistically determined based on the simulated half-life and end point. Transcripts with end points beyond g were terminated at g (or g + d for alternative cleavage site simulations), but the longer end points affected the probability of cleavage since they represent transcripts that were completed before the end of the labeling period. Mean library preparation insert sizes and standard deviations were used during the in-silico fragmentation and size-selection steps and the first 50nt of the fragments were used as simulated reads.

### Estimating 3’ end cleavage half-lives in *Drosophila melanogaster* S2 cells

#### Filtering cleavage sites

To identify cleavage sites for which we could confidently estimate cleavage rates, we conditioned on 4 criteria: (1) sites in genes with TPM > 5 in the total RNA libraries; (2) sites whose use as a polyA site is supported by 3p-seq data (see above); (3) sites that do not overlap introns (defined using RNA-seq junction reads), since splicing efficiency over the 4sU time course might confound measurement of reads supporting uncleaved transcripts; (4) sites for which the associated total read region (**Fig 1C**) does not overlap an intron, since splicing efficiency might similarly confound total transcript measurements and (5) sites with at least 1 uncleaved or cleaved read in any time point. Finally, alternative cleavage sites within 50 nt of each other were filtered out. Using these criteria, we retain 2, 857 constitutively used (PAU > 0.95) and 603 alternatively used (PAU > 0.05 & PAU < 0.95) sites.

#### Processing RNAPII ChIP-seq data and estimating length of readthrough region

To define a readthrough region, defined as the region that RNAPII continues to travel after cleavage, we used publicly available RNAPII ChIP-seq data from *Drosophila melanogaster* S2 cells. Specifically, fastq files from GSE112608 (61) were downloaded from GEO and mapped using STAR as described above. To estimate the length of the readthrough region, we used DoGFinder (62), run with default parameters and a conditioning on a minimum downstream of gene (DoG) length of 100 nt.

#### Extracting informative reads and modeling cleavage half-lives

For each of the cleavage sites retained for further analyses, informative reads for cleavage site and total region were counted using bedtools. For cleavage site reads, we required at least a 10nt overlap with both terminal exon (upstream of cleavage site) and readthrough region (downstream) for cleavage site reads. We defined the total region as the region located (2[*insert length*] + 2[*library std dev*]) upstream of the cleavage site and counted all reads that overlapped this region. Reads were combined across the three replicates from each labeling period to increase power to model half-lives and half-lives were estimated using a constant Drosophila transcription rate of 1500 nt/min (63, 64).

### Nucleotide Composition and kmer analysis

To obtain nucleotide composition information, genome sequence was extracted around each cleavage site using *bedtools getFasta -s*. The proportion of each nucleotide (A, C, G, U) present across all sites or relevant sites categorized by half-life measurements was calculated for each nucleotide position in the specified windows. The same genome sequences were scanned for the 10 most prevalent polyadenylation signal (PAS) kmers specified in Sanfillipo *et al.* (28) using a custom python script to identify the position and identity of PASs around each cleavage site.

## RESULTS

The processing of RNA molecules cleavage is thought to occur co-transcriptionally, with molecular steps initiating rapidly after recognition of splice sites or PASs for mRNA splicing and 3’ end cleavage, respectively. Thus, to measure the kinetics of RNA events, it is necessary to profile transient nascent RNA intermediates at the proper timescale. This is challenging to achieve with single locus quantitative PCR or single molecule imaging techniques, which are limited in either quantitative or temporal resolution, respectively, for rapid RNA processing mechanisms. To overcome these challenges, we recently developed a modeling approach to estimate the rates of mRNA splicing using data from high-throughput sequencing of nascent RNA labeled with 4-thio-uridine (4sU) over short labeling periods (24). Specifically, our approach uses the ratio of spliced exon-exon junction reads to unspliced intron-exon junction reads to model the half-life of intron excision. Since the progressive incorporation of 4sU over the labeling period results in a distribution of times since completion of synthesis of an intron - and thus of lifetimes over which an intron could be spliced - our model explicitly integrates over possible polymerase locations that could contribute junction reads to estimate the probability of observing a junction ratio consistent with a given half-life. Since progressive, short time period metabolic labeling of nascent RNA also provides sufficient read coverage at the 3’ end of a gene, we set out to adapt our splicing rates model to estimate the rates of 3’ end pre-mRNA cleavage.

### Identifying signatures of 3’ end mRNA cleavage in nascent RNA

We previously estimated mRNA splicing rates by empirically measuring exon-exon and intron-exon junction reads from 4sU labeled nascent RNA-seq data, which unambiguously quantify spliced and unspliced molecules, respectively. To adapt this approach to estimate rates of 3’ end cleavage, we must first identify nascent RNA reads informative about cleaved and uncleaved molecules. Reads that directly overlap the cleavage site (CS) unambiguously arise from uncleaved molecules, however, there are no such reads that are exclusive to cleaved molecules. To identify a population of reads that could be used to quantify cleaved molecules, we used our previously published nascent RNA sequencing dataset in *Drosophila melanogaster* S2 cells, which includes 5, 10, and 20 minute 4sU labeling time points, to assess patterns of read distribution around the CS. We find a depletion of nascent RNA reads around annotated CS, with the extent of read depletion growing more pronounced as labeling time increases (**FIGURE 1A**). We hypothesized that this temporally associated depletion reflects increasing numbers of cleaved molecules, which cannot contribute reads directly over the CS. Indeed, similar patterns of temporally associated read depletion were observed in reads generated from a simulated 4sU-labeled nascent sequencing dataset **(FIGURE S1, FIGURE S2A)**.

We rationalized that we could obtain an expected value for the number of cleaved reads by using reads upstream of the CS to initially estimate the total number of transcripts that could be cleaved. Molecules resulting from RNAPII readthrough downstream of the CS are less informative since they are rapidly degraded by XRN2, reflected in the decrease in 4sU-seq reads in the region downstream of the CS (**FIGURE 1A**). Despite the biological differences in the lifespan of upstream mRNA and downstream readthrough molecules, the shape of the dip in read distribution was symmetrical around the CS. This pattern was consistent with RNA-seq edge effects, where size selection for narrow fragment length distributions during library preparation causes a depletion of reads at the ends of molecules. Cleavage of nascent transcripts would result in edge effects for both the upstream and downstream molecules. Read coverage in the simulated data confirmed that the width of the read depletion distribution is associated with library preparation fragment length after size selection (**FIGURE 1B**). Thus, we realized that it is necessary to control for the average fragment length in a library when identifying a region from which to quantify the total number of cleavable transcripts.

Together, these insights across the in-vivo and simulated 4sU-seq datasets led us define two sets of cleavage-informative reads: (1) reads overlapping the CS to quantify uncleaved transcripts (uncleaved reads) and (2) reads overlapping a region two fragment lengths upstream of the CS to quantify cleavable transcripts (total reads) and estimate the expected number of cleaved transcripts (cleaved reads; Methods, **FIGURE 1C**). Reassuringly, we see that the ratio of uncleaved reads to total reads decreased over 4sU labeling time in the 4sU-seq data from fly S2 cells, consistent with more molecules being cleaved over time (**FIGURE S2B**). While this ratio correlates with cleavage half-lives in the simulated nascent data, the strength of the correlation is dependent on the length of the readthrough region (**FIGURE S2C**). This suggests that robust half-life estimation must account for polymerases continuing to contribute labeled reads while transcribing in the readthrough region, just as we accounted for the regions downstream of the 3’ splice site in our previous approach for estimating splicing rates.

### Estimating the rate of 3’ end cleavage in fly S2 cells

To estimate the half-life of 3’ end cleavage, we modified our splicing rate model to model the probability of observing an given ratio of uncleaved to cleaved reads (estimated empirically) integrating over a region accounting for both polymerase readthrough region (estimated using ChIP-seq data; **FIGURE S2D**) and the labeling time (Methods). The model (**FIGURE 1C**) makes three major assumptions: (1) a constant transcription rate over the entire region past the 3’ CS, (2) RNAPII localization on the DNA is indicative of actively transcribing RNAPII after cleavage, and (3) cleavage events will always occur to completion, thus we can only estimate the rates of constitutively used cleavage sites. This model was able to accurately assign absolute rates of 3’ end cleavage to each CS in our simulated 4sU-seq data (Spearman *rho* = 0.98; **FIGURE 1D**), across a range of simulated half-lives, expression levels, library preparation conditions, and genomic parameters. Furthermore, we see no detectable bias in the relative accuracy of half-life estimation across the simulated distributions of expression levels (**FIGURE S2E**), readthrough regions (**FIGURE S2F**), and transcription elongation rates (**FIGURE S2G**). Overall, results from our simulations indicate that modeling of progressive labeling 4sU-seq data can be used to robustly estimate rates of 3’ end cleavage.

As stated above, one of the limitations of our initial estimation of cleavage-informative reads is that our model assumes all reads are derived from transcripts that are constitutively cleaved at a site, which does not hold for alternatively used cleavage sites(5% > PAU < 95%; **FIGURE S3A**). The presence of reads from transcripts that are cleaved at an alternative site within the same gene results in an artifactual increase in uncleaved reads and an overestimation of 3’ end cleavage half-lives for alternative sites in both simulated (**FIGURE 1E**, **FIGURE S3B**) and 4sU-seq data (**FIGURE S3C**). Thus, we modified our read estimation scheme to proportionally allocate informative reads to alternative cleavage sites based on (1) the relative usage of the sites, (2) genomic order and position of the sites, and (3) the first two assumptions used for estimating constitutive rates above (Methods; **FIGURE S4**). Our 3’ end cleavage rates model performs quite well to estimate the 3’ end cleavage half-lives of alternative sites using these adjusted reads (**FIGURE 1F**; **FIGURE S3D**).

### Alternative 3’ cleavage sites are cleaved slower than constitutive sites

We applied this novel model for estimating 3’ end cleavage rates to our previously published time course of 4sU-seq data from *Drosophila melanogaster* S2 cells (24). After filtering for sites with sufficient power to estimate rates (Methods), we modeled cleavage half-lives for 2, 857 constitutive (PAU > 95%) and 603 alternative (5% < PAU < 95%) sites in S2 cells (**FIGURE 3A**). Within this set, we were unable to obtain half-lives for 185 constitutive and 354 alternative sites (after read adjustment) that had no uncleaved reads at any time-point, suggesting that these sites are cleaved faster than the resolution afforded by our labeling time-course. Additionally, we were only able to estimate rates of gene-proximal (upstream most) alternative sites, likely due a combination of reduced usage of distal sites and the 4sU-seq 5’ bias leading to decreased read coverage at sites further from the transcription start site. Among the remaining sites, the median cleavage half-life (t_1/2_) was 35.3 and 43.3 seconds (standard deviation of 0.03 and 120 seconds) for constitutive and alternative sites, respectively (**FIGURE 2A**).

**Figure 2.**
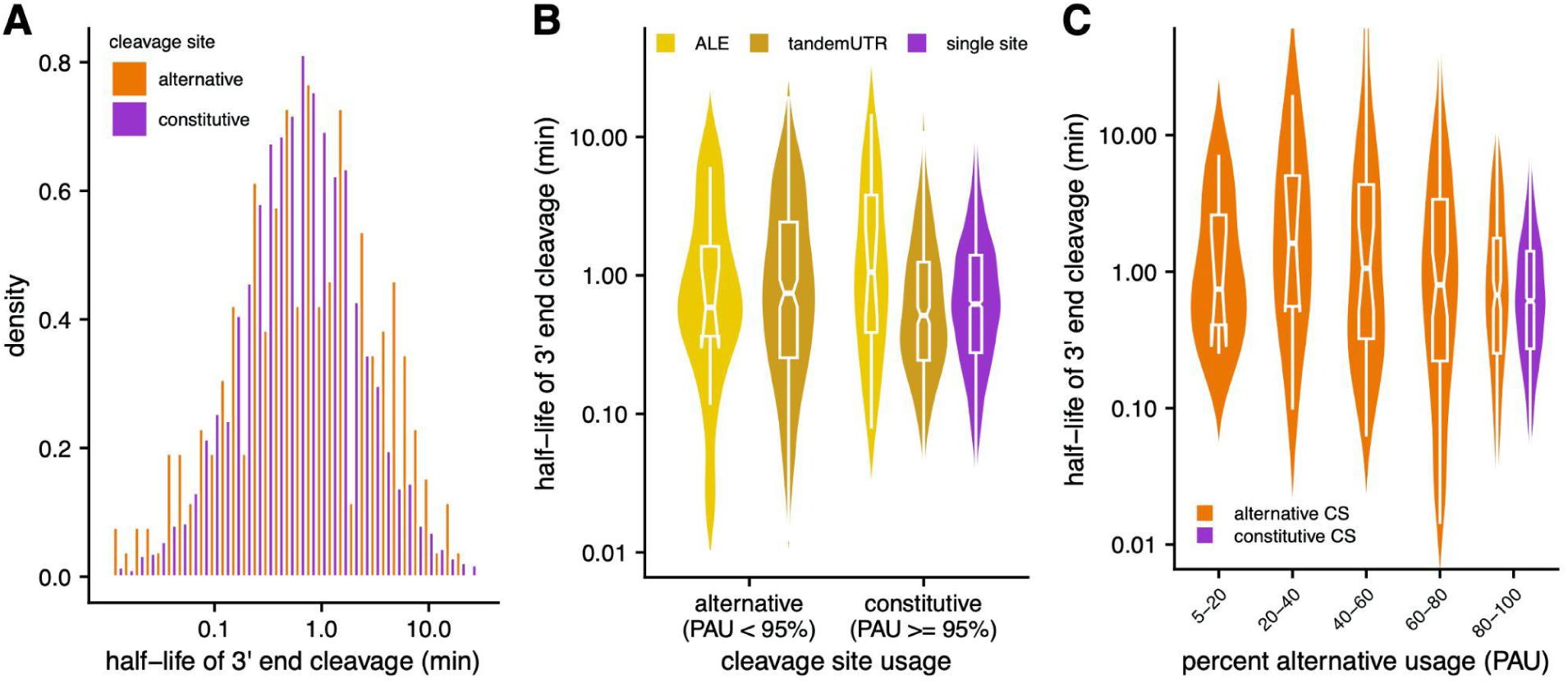
Estimating cleavage rates at constitutive and alternative sites. **(A)** Distribution of 3’ end cleavage half-lives for constitutively (PAU > 95%; *purple*) and alternatively cleaved sites(5% < PAU < 95%; *orange*). **(B)** Distribution of estimated 3’end cleavage half-lives separated by annotated site usage. ALE refers to alternative last exons (alternative n = 12; constitutive n = 24), tandemUTR refers to tandem 3’ UTRs where more than one cleavage site are in the same exon (alternative n = 251; constitutive n = 359), and single sites are in genes with only one annotated cleavage (constitutive n = 2, 398) site. **(C)** Distribution of estimated 3’end cleavage half-lives for quintiles of PAU values (n = 8, 8, 31, 61, and 155 for alternative sites across quintiles from 5-20% to 80-100%), separated by alternatively used (*orange*) and constitutively used cleavage sites (*purple*).

**Figure 3.**
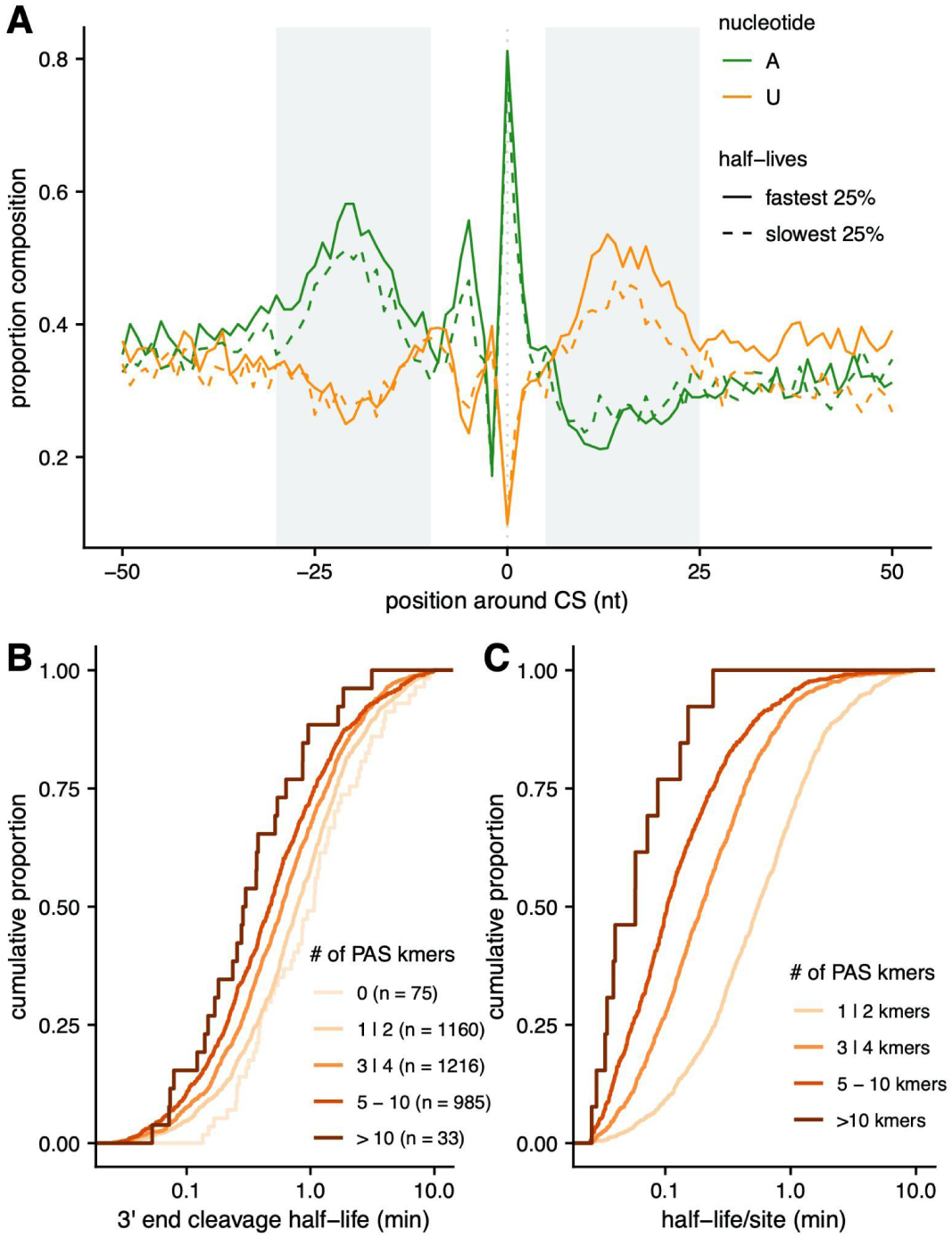
RNA sequence elements are associated with 3’ end cleavage half-lives. **(A)** The proportion of cleavage sites (*y-axis*) with adenine (*green*) or uridine (*orange*) nucleotides at each position in the 100nt (*x-axis*) surrounding cleavage sites with the fastest (*solid line*) and slowest (*dashed line*) 25% half-lives. **(B)** The cumulative distribution of cleavage half-lives (*x-axis*) for sites grouped by the number of PAS kmers identified in the 200nt around each cleavage site (*colored lines*). **(C)** The cumulative distribution of cleavage half-life divided by the number of PAS kmers identified in the 200nt around each site.

On average, half-lives for alternative cleavage sites were slightly but significantly slower than constitutive sites (Welch’s two sample test p-value = 0.0006). Similarly, sites that are associated with alternative last exons trend towards being cleaved slower on average (**FIGURE 2B**), despite being constitutively used in S2 cells, suggesting a small delay in recognizing or initiating cleavage at alternative sites that may be temporally coupled with local splicing decisions. Finally, the 3’ end cleavage half-lives of alternative sites were weakly associated with the relative usage of the sites (**FIGURE 2C**), such that sites with less relative usage were cleaved slower. Together, these observations suggest that the kinetics of 3’ end cleavage may play a role in the regulation of alternative cleavage decisions.

### Faster cleavage is associated with the strength and number of PAS elements

The initiation of 3’ end cleavage is mediated by recognition of multiple cis-regulatory RNA elements located both upstream and downstream of the CS. These include the polyadenylation signal and downstream sequence element (DSE), positioned 10-30nt upstream and ∼10-40nt downstream of the CS, respectively (1, 28–30). To understand how cis-elements influence the rate of 3’ end cleavage, we first calculated nucleotide composition in the 100nt window around constitutive cleavage sites. We broadly recapitulate the known nucleotide distributions across all sites, characterized by (1) an A at the CS, (2) an overall enrichment of AU content around the CS, (3) an A-rich region 10-30nt upstream of the CS, and (4) a U-rich region 5-25nt downstream of the CS (**FIGURE S5A**, (1, 28–31)). Strikingly, we see that these sequence characteristics are enhanced in the cleavage sites with the fastest half-lives, combining across both constitutive and alternative sites, with a greater proportions of A and U content upstream and downstream of the CS, respectively (**FIGURE 3A**).

Given the known importance of PAS strength in CS usage (2, 23), we next turned to specifically investigate how the sequence and position of the PAS influences the rate of 3’ end cleavage. To do so, we identified positions for the PASs predominantly used in Drosophila (28) by scanning for the 10 mostly commonly used 6-mers in the 200 nucleotides upstream of sites for which we measured cleavage rates. Consistent with previous observations (28), we observe that for 60% of genes, the closest PAS 6-mer is within 10-30nt upstream of the CS and 66% of these 6-mers match the strongest AAUAAA, AUUAAA, or AAUAUA PAS motifs thought to be present at the majority of fly genes (28) (**FIGURE S5B**). We do not see any significant associations between cleavage half-lives and either the position (**FIGURE S5C**) or the sequence of the closest PAS kmer (**FIGURE S5D**). Instead, we find that cleavage sites with a greater number of PAS kmers in the 200nt upstream of the CS have faster cleavage half-lives (**FIGURE 3B**). This could indicate that PASs act cooperatively, where each additional element increases the probability of CPA machinery binding in an additive fashion. To evaluate this possibility, we calculated the half-life per kmer and and again see the same relationship, where cleavage sites that have more kmers are being cleaved faster per sequence element (**FIGURE 3C**). This observation suggests a non-linear multiplicative relationship where multiple elements cooperate to increase the rate of cleavage more than the rate driven by any individual site.

### 3’ end cleavage rates are associated with local RNA polymerase II dynamics

The rate of cleavage has the potential to be associated with RNAPII localization, since PAS recognition likely occurs while the nascent RNA is emerging from the RNAPII exit channel. Since 3’ end cleavage initiates the decoupling of the mRNA molecule from actively elongating RNAPII, we hypothesized that cleavage kinetics may be influenced by RNAPII dynamics downstream of the CS. Consistently, we see that the rate of 3’ end cleavage is significantly associated with the length of the readthrough region, or the region that RNAPII transcribes after the CS (Spearman ⍴ = 0.34, p-value < 2.2 × 10^-16^; **FIGURE 4A**). We do not observe this association in simulated data, indicating that this observation is not due to biases in our modeling approach. The observation that cleavage sites with faster half-lives were associated with shorter readthrough regions suggests that RNAPII dynamics may be limited in some fashion in the region after the CS, as was recently suggested to occur in yeast and human cells (32, 33). To investigate this, we used RNAPII ChIP-seq data from Drosophila S2 cells to look at RNAPII occupancy around cleavage sites. For these analyses, we only used constitutive sites to avoid overlapping signals from alternative sites in the same genes. As has been observed previously, we observe decreased RNAPII occupancy just upstream of the CS and an immediate increase in occupancy downstream of the CS, regardless of cleavage rate (**FIGURE 4B**). However, this increase in RNAPII occupancy is more pronounced in the first 250nt downstream of the CS for sites with the shortest half-lives.

**Figure 4.**
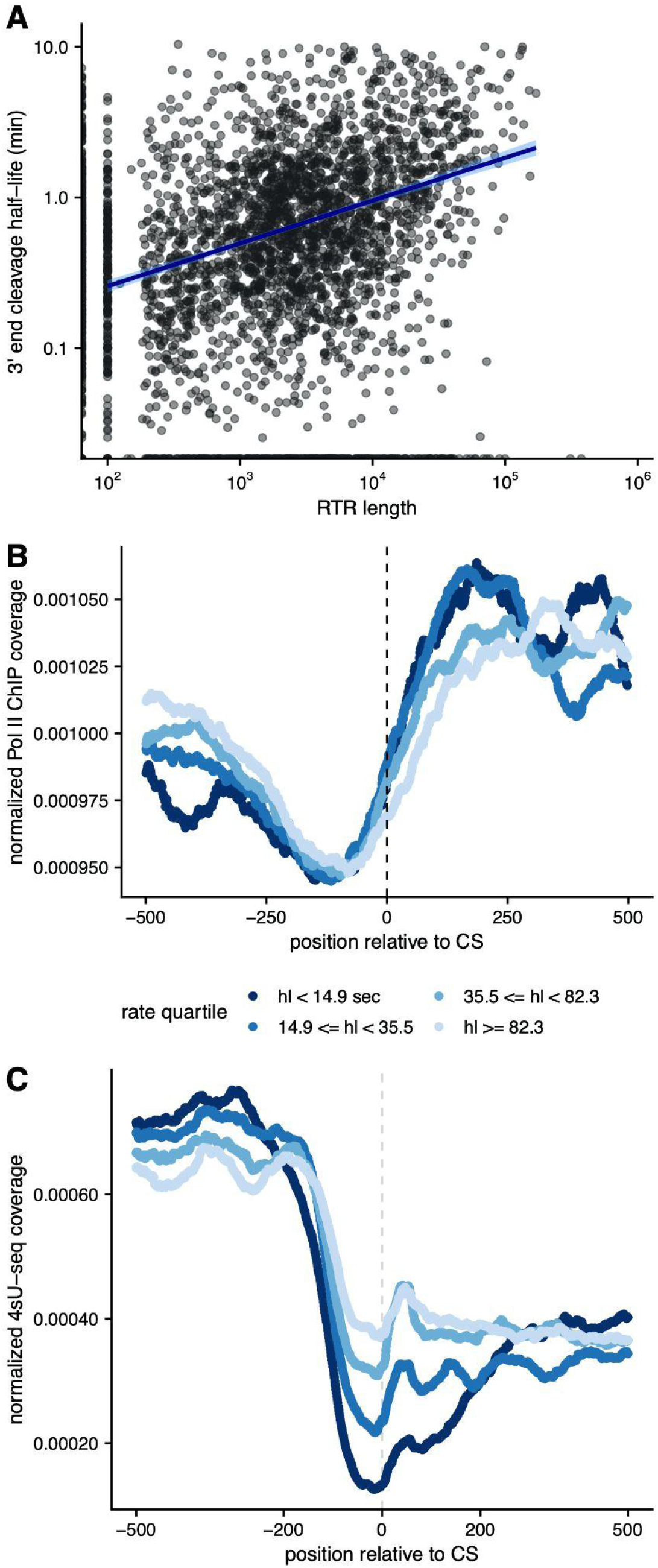
3’ end cleavage half-lives are associated with RNA polymerase dynamics at the 3’ end of the gene. **(A)** Cleavage half-lives (*y-axis*) *versus* the lengths of readthrough regions(in kb, *x-axis*), with a linear fit (*blue*). **(B)** Meta-gene plot of normalized RNA PollII ChIP-seq read coverage (*y-axis*) in a 1000nt region centered on constitutively used cleavage sites (*x-axis*), with sites separated by quartiles of cleavage half-lives (*blue lines*). **(C)** Meta-gene plot of normalized 4sU-seq read coverage for the 5m 4sU labeling time point (*y-axis*) in a 1000nt region centered on constitutively used cleavage sites (*x-axis*), with sites separted by quartiles of cleavage half-lives (*blue lines*).

Together, these observations indicate a relationship between cleavage rates and RNAPII dynamics in the 250nt downstream of the cleavage site. Specifically, this observation suggests that RNAPII pausing or a reduction in RNAPII speed may underlie faster cleavage. Concurrently, faster 3’ end cleavage may have consequences on RNAPII processivity, since the earlier appearance of an unprotected 5’ end on the readthrough transcript may result in faster XRN2-mediated degradation of the readthrough transcript. This could limit the distance that RNAPII is able to transcribe before XRN2 catches up and promotes transcription termination. Consistent with this hypothesis, we see fewer readthrough transcripts for cleavage sites with faster half-lives, especially in early nascent data when readthrough transcripts are less likely to be mostly fully degraded (**FIGURE 4C, FIGURE S6**).

## DISCUSSION

Recent development of high-throughput sequencing techniques that capture nascent RNA over time has made genome-wide kinetic profiling of RNA maturation possible. Though rates of mRNA splicing have been estimated globally, the rate at which an mRNA is cleaved and polyadenylated to complete the maturation process has never been investigated. In this study, we introduced a novel method to measure the rates of site-specific 3’ pre-mRNA cleavage. We applied this approach to estimate the half-lives of 3’ end cleavage in *Drosophila melanogaster* S2 cells and examine genomic features that may account for variability in cleavage rates.

We find that, on average, cleavage at sites that are alternatively cleaved occurs 23% slower than at constitutive cleavage sites. Furthermore, constitutively used cleavage sites in genes known to have alternative sites are also cleaved slower than sites in genes with only one possible CS. Together, these observations suggest that capacity for alternative site usage in a gene may lead to delays in 3’ end cleavage and polyadenylation, perhaps until all cleavage sites have been transcribed. However, after adjusting informative read counts by the proportional CS usage, we were only able to estimate cleavage half-lives for the gene proximal-most sites in any gene. While we cannot discount the possibility that cleavage at distal sites is often too fast to estimate, our inability to estimate half-lives at these sites is likely due to the inherent 5’ sequencing bias particularly present in early 4sU labeling timepoints. Thus, our read adjustment approach for counting informative reads may be especially underpowered at distal sites and higher sequencing depth is necessary to estimate the rates of cleavage at these sites.

Sites with faster cleavage half-lives also tend to have enrichments of sequence features traditionally associated with more efficient 3’ end cleavage and polyadenylation - namely, an A-rich region upstream of the CS, a U-rich downstream element distal to the CS, and a higher density of kmers associated with polyadenylation signals. These observations suggest that the speed of cleavage could be regulated by the recruitment of regulatory features. However, our analyses suggest that PAS motifs do not act independently but rather cooperate such that the presence of multiple motifs increases the rate of cleavage more than expected for the addition of a site. This is consistent with reports that genes can have clusters of efficiently cleaved, closely spaced 3’ isoform endpoints whose usage is associated with temporal regulation of RNA processing (33). One possibility is a model in which the local density of these motifs is the determining feature for increasing the concentration of cleavage and polyadenylation machinery near the cleavage site and hastening the cleavage reaction.

The presence of G-rich sequences - putatively enriched with G-quadruplexes - downstream of cleavage sites has been previously posited to be associated with enhanced cleavage and polyadenylation efficiency by aiding in the recruitment of necessary regulatory factors or promoting an open structural conformation of all the elements (29, 34–36). Furthermore, G-rich or G-quadruplex containing regions have also been hypothesized to function as roadblocks or pause sites for RNAPII elongation, serving to slow down or pause RNAPII downstream of cleavage sites and facilitate rapid cleavage (33, 35, 37). Giesberg *et al.* recently found that slower RNAPII speed at downstream of cleavage sites promotes cleavage and polyadenylation and the speed of RNAPII is mediated by GC-content in this region (33). Consistent with this observation, we do observe evidence for increased RNAPII dwelltime downstream of cleavage sites with the fastest half-lives, but do not observe a relationship between GC content and and cleavage rates. Our observations support the hypothesis that slower transcription elongation enables faster 3’ end cleavage, but that the relationship between cleavage speed and RNAPII dynamics is not mediated by G-containing sequence elements. It is also possible that both the speed of cleavage and RNAPII are independently linked to increased polyA site usage in steady-state mRNA.

Alternatively, the increase in RNAPII density downstream of the fastest cleavage sites might be due to RNAPII stalling during termination and chromatin disassociation (38). The torpedo model posits that termination is initiated by XRN2, which performed exonucleolytic 5’-3’ degradation of the readthrough transcript, catching up to the polymerase and promoting dissociation from DNA (38, 39). Faster cleavage likely allows XRN2 to gain a kinetic advantage (40), allowing XRN2 to interact with RNAPII before it has a chance to advance far downstream of the CS. We observe fewer reads in the readthrough region downstream of the fastest cleaved sites, consistent with the idea that XRN2 degradation of the downstream molecule begins faster. Future studies to measure the rate of XRN2 exonucleolytic degradation itself would shed light on the relationship between the dynamics of 3’ end cleavage and polymerase and sequence of events occurring during transcription termination.

More broadly, our approach to estimate the rate of 3’ end cleavage can help to provide insights into the spatio-temporal sequence of events necessary to create an mRNA molecule. When leveraged in tandem with other nascent RNA modeling approaches to measure transcription elongation and splicing rates, these tools together can be used to elucidate the rate-limiting steps for mRNA biogenesis across biological contexts. Studies have found increasing evidence of the co-regulation of splicing and cleavage on the same molecule, with evidence that inefficient splicing is associated with inefficient 3’ end cleavage and increased readthrough transcripts in both yeast and human cells (41, 42). These observations support potential kinetic coupling between splicing and cleavage events. Similarly, recent evidence that alternative last exons are likely regulated by cleavage rather than splicing mechanisms (43) suggests that the timing of cleavage might compete with or dictate the timing of splicing for these alternative decisions, as suggested by window of opportunity models (20).

Alternative cleavage and polyadenylation is known to be tightly regulated in many biological contexts, including neuronal development (44–49), circadian rhythms (50, 51), the immune response (19, 52, 53), and many cancer subtypes (54, 55). In each of these contexts, the timing of regulatory events matters to ensure proper cellular differentiation, clock timing, survival from stimulation stress, or proliferative advantages, respectively. Thus the kinetics of cleavage decisions may specifically be regulated across these and other contexts to ensure appropriate 3’ UTR selection. For instance, a kinetic balance between splicing and cleavage was shown to be perturbed by a cancer-associated mutation in the U2AF1 splicing factor, leading cleavage to no longer be rate-limiting for any molecules of the B-globin gene (27) and broadly suggesting that this commonly occurring mutation may be selected to influence kinetic competition in human cancer cells. Applying our method to measure the rates of 3’ end cleavage in a diversity of cell types and biological system would enable further understanding how widespread kinetic co-regulation or competition is and how the timing of regulatory events may dictate context-specific isoform usage.

## AUTHOR CONTRIBUTIONS

## ACKNOWLEDGEMENTS

We thank Chris Burge and Anish Shah for initial discussions on developing the method, Elisa Donnard and Manuel Garber for suggesting the use of RNAPII ChIP-seq data to estimate RTR lengths, and the Pai Lab for helpful discussions and feedback on the manuscript.

