## Supplementary Material for "Genome-wide kinetic profiling of pre-mRNA 3’ end cleavage"

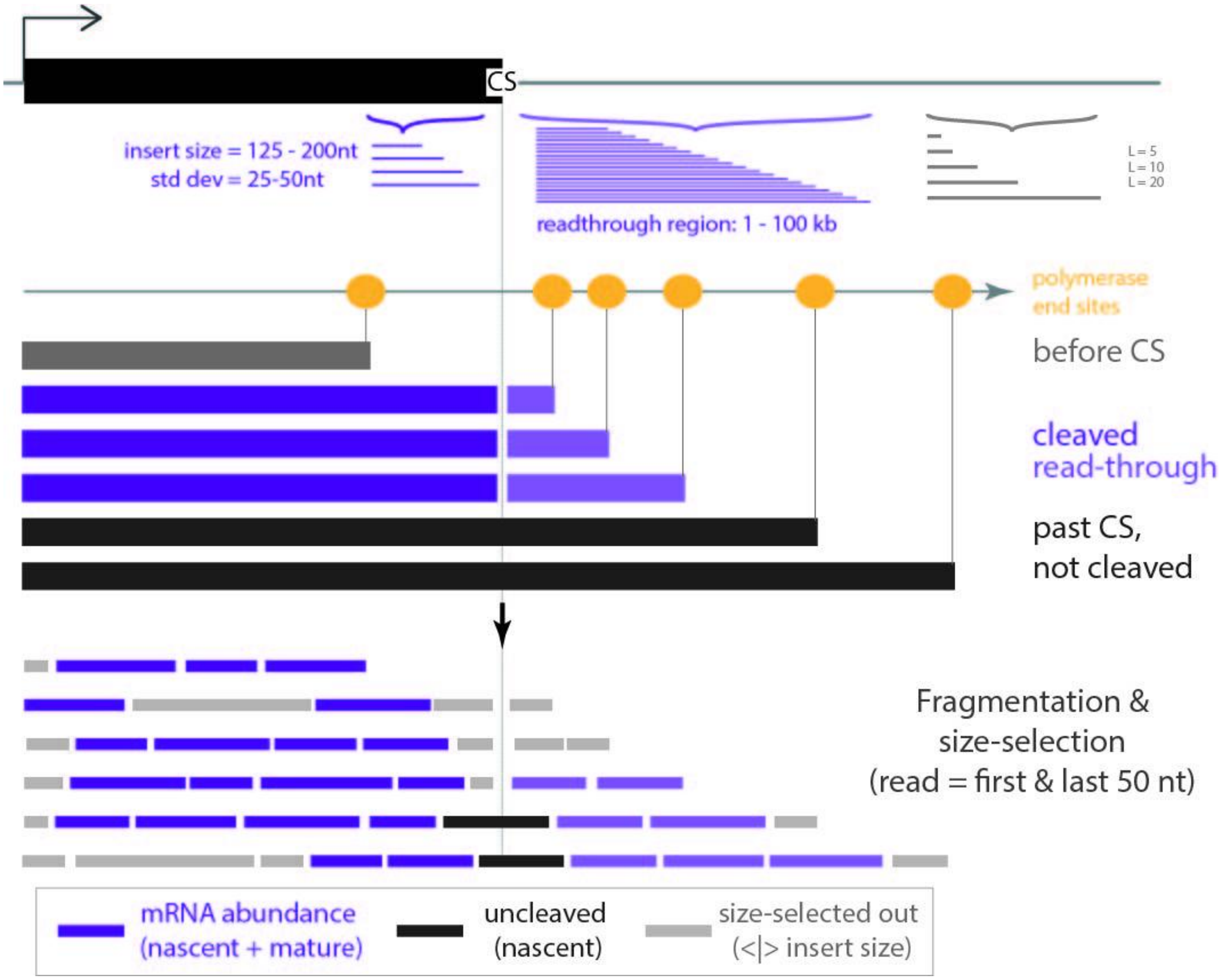


**Figure S1. Schematic of read simulations around cleavage sites.** 4sU-seq reads were simulated across a range of parameters (see Methods). Transcripts can be separated by 3 categories: (1) not yet transcribed past cleavage site (*top*, *grey*), (2) transcribed past the cleavage site and cleaved (*middle*, *purple*), and (3) transcribed past the cleavage site but not yet cleaved (*bottom*, *black*). During the library preparation fragmentation and size-selection steps, fragments that are longer or shorter than the library insert size are selected out (*grey fragments*).

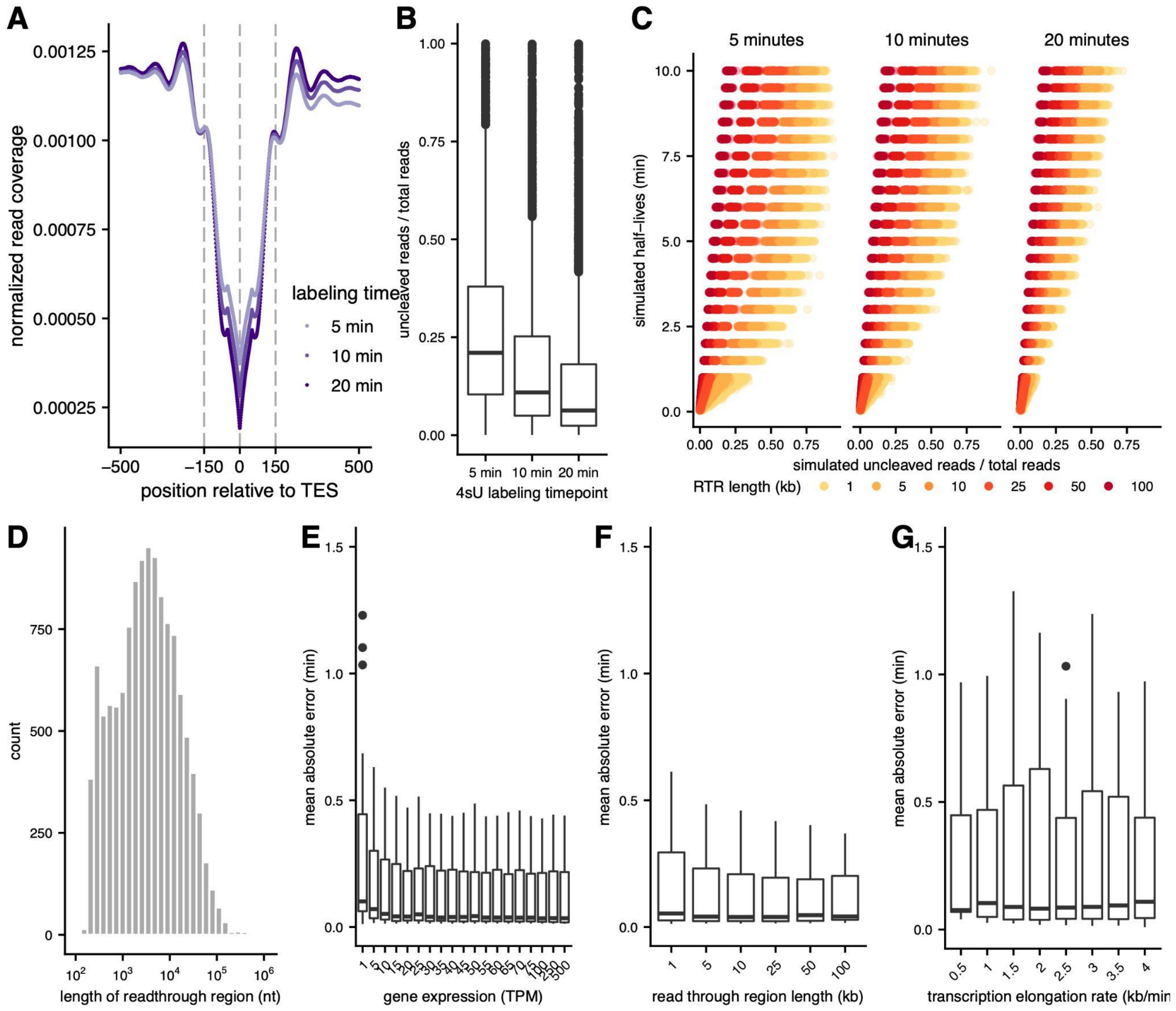


**Figure S2. Testing a mathematical model to estimate site-specific rates of 3' end cleavage. (A)** Meta-gene plot showing average simulated read coverage (*y-axis*) in a 1000nt region centered on simulated 3' end cleavage sites (*x-axis*), separated by simulated labeling timepoints (*purple lines*). The dashed line indicates the cleavage site (coordinate 0), while dotted lines indicate the simulated library fragment length (150nt). **(B)** Ratio of uncleaved to total reads for cleavage sites for nascent RNA collected 5 min, 10 min, 20 min after 4sU labeling. The overall decrease in uncleaved to total reads over time indicates increased completed cleavage. **(C)** Simulated cleavage half-lives (y-axis) versus the ratio of uncleaved to total reads in simulated data (y-axis) across labeling time points. The correlation between these metrics is biased by the simulated length of the readthrough region (RTR; colors). **(D)** Distribution of read-through regions estimated using PolII ChIP-seq data. **(E-G)** Mean absolute error in the estimation of cleavage half-lives across a range of simulated half-lives (y-axis) for a range of gene expression values (simulated as TPM; *x-axis*; **E**), readthrough region lengths (simulated in kilobases, *x-axis*, **F**), and transcription elongation rates (simulated in kb/min, *x-axis*, **G**).

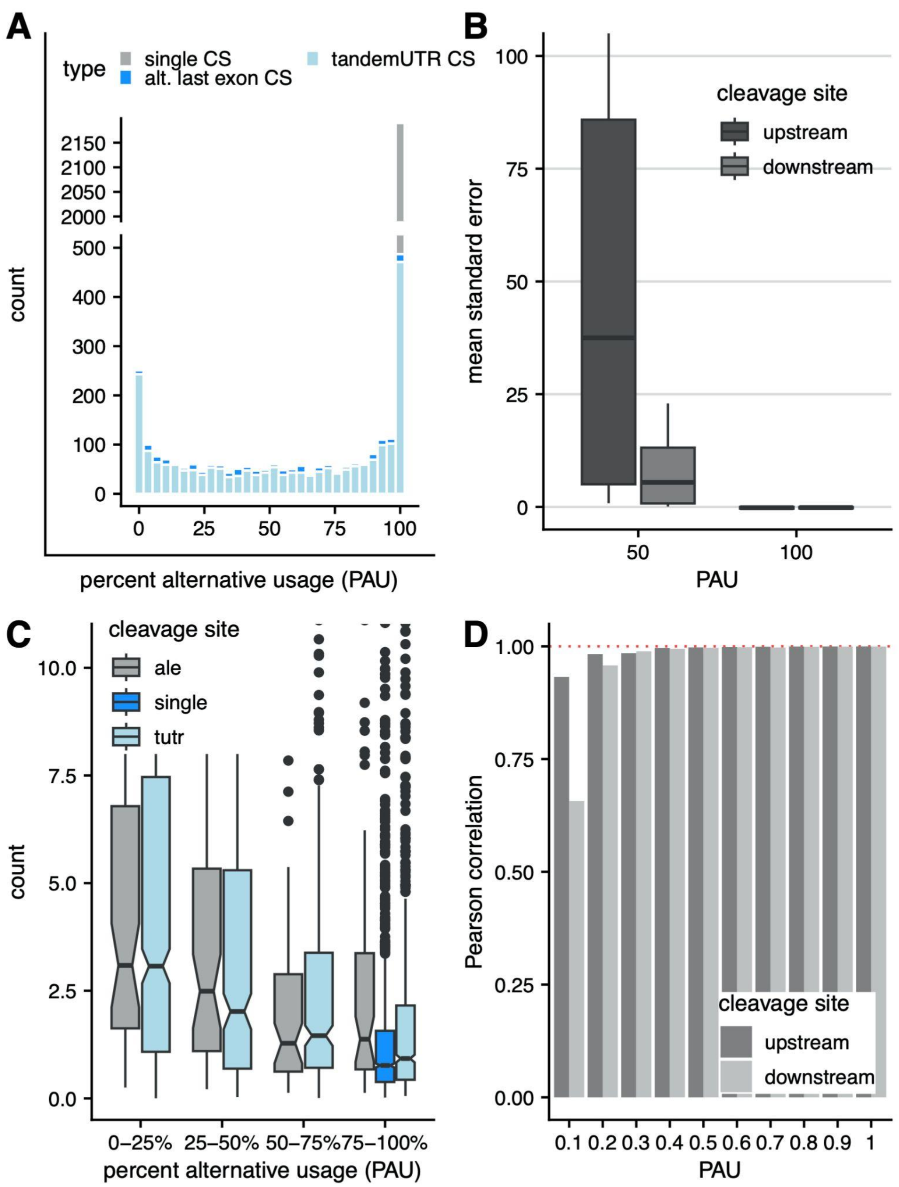


**Figure S3. Testing approaches to estimate rates of 3' end cleavage at alternative sites. (A)** Distribution of PolyA Usage (PAU) values, separated by annotated alternative exon type (colors). **(B)** Mean standard error in the estimation of cleavage half-lives across a range of simulated half-lives for upstream or downstream alternative cleavage sites with a PAU of either 50% or 100%. Uncorrected read counts do well at modeling the cleavage half-life when considering a PAU = 100%, but overestimate cleavage half-lives with PAU = 50%. **(C)** Estimation of cleavage half-lives (y-axis) across *Drosophila* cleavage sites with variable PAU values (x-axis) with uncorrected read counts. **(D)** Pearson correlations between the estimated and simulated cleavage half-lives across a range of simulated half-lives (*y-axis*) for upstream and downstream alternative sites, across a range of PAU values for each. The read correction performs well for sites with PAU > 0.3.

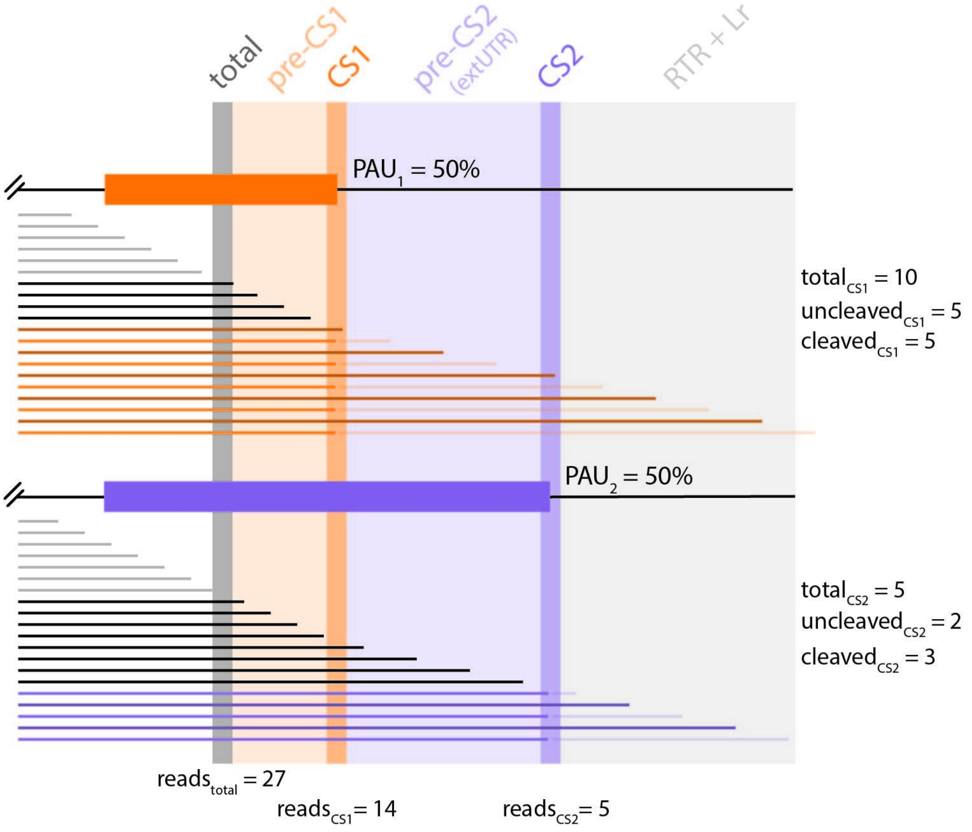


pre-CS1 proportion = (2* insert + 2*insert$_{StdDev}$) / (2* insert + 2*insert$_{StdDev}$ + extUTR + RTR + Lr)

total$_{CS1}$ count = (reads$_{total}$ * PAU$_1$) - (reads$_{total}$ * pre-CS1 proportion)

uncleaved$_{CS1}$ count = reads$_{CS1}$ - (reads$_{total}$ * (1 - PAU$_1$))

cleaved$_{CS1}$ count = CS1 total count - CS1 uncleaved count

pre-CS2 proportion = extUTR / (extUTR + RTR + Lr)

total$_{CS2}$ count = (reads$_{total}$ * PAU$_2$) - (reads$_{total}$ * pre-CS1 proportion) - (reads$_{total}$ * pre-CS2 proportion)

uncleaved$_{CS2}$ count = reads$_{CS2}$ - (uncleaved$_{CS1}$ count - (uncleaved$_{CS1}$ count * pre-CS2 proportion)

cleaved$_{CS2}$ count = CS2 total count - CS2 uncleaved count

**Figure S4. Schematic of read adjustment for estimating rates of alternative cleavage sites.** 4sU-seq reads were generated across a range of parameters for a simulated gene with two alternative cleavage sites (see Methods). Transcripts for each site can be separated by 4 categories: (1) not yet transcribed past the region from which the total number of transcripts is counted (*top*, *grey*), (2) transcribed past the total region but not yet transcribed to the cleavage (*middle top*, *black*), (3) transcribed past the cleavage site but not yet (*middle bottom, dark orange and dark purple*), and (4) transcribed past the cleavage site and cleaved (*bottom*, *orange and purple*). To account for reads specifically informative for each of the cleavage sites, reads are weighted by the site-specific PAUs and adjusted to subtract reads arising from transcripts that cannot or will not be cleaved at a given site (*bottom*, Methods).

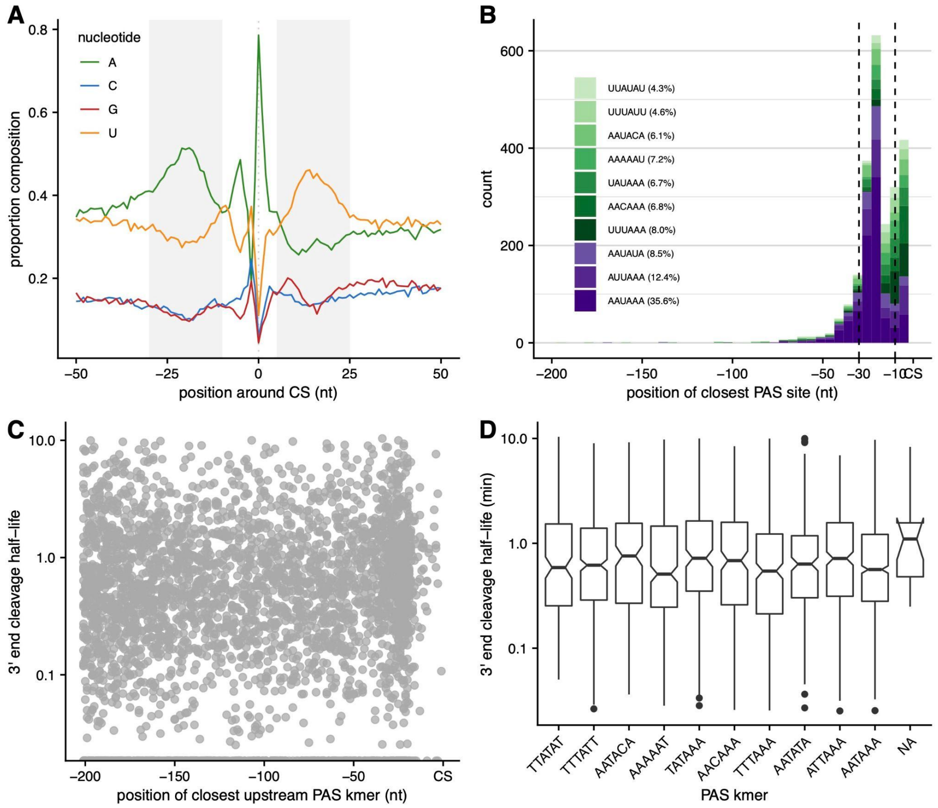


**Figure S5. Identifying RNA sequence elements near 3' end cleavage sites. (A)** The proportion of cleavage sites (*y-axis*) with each of the four nucleotides (*colored lines*) in the 100nt around cleavage sites (*x-axis*). **(B)** The distribution of PAS hexamers identified in the 200nt upstream cleavage sites (*x-axis*), separated by the 10 most commonly used PAS kmers (*purple* for 3 major kmers and *green* for minor kmers). **(C)** Cleavage half-life (*y-axis*) *versus* position of closest PAS kmer (*x-axis*). **(D)** The distribution of cleavage half-lives (*y-axis*) for the 10 most commonly used PAS kmers (*x-axis*)**.**

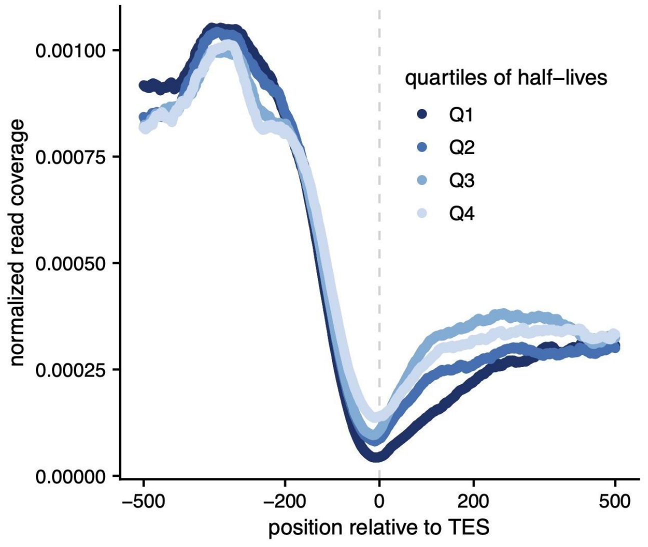


**Figure S6.** Meta-gene of normalized total RNA-seq read coverage (*y-axis*) in a 1000nt region centered on constitutively used cleavage sites (*x-axis*) with sites separated by quartiles of cleavage half-lives (*blue lines*).
